# Methylation by multiple type I restriction modification systems avoids influencing gene regulation in uropathogenic *Escherichia coli*

**DOI:** 10.1101/2021.01.08.425850

**Authors:** Kurosh S. Mehershahi, Swaine L. Chen

**Author notes:** Corresponding author. (SLC).

## Abstract

DNA methylation is a common epigenetic mark that influences transcriptional regulation, and therefore cellular phenotype, across all domains of life, extending also to bacterial virulence. Both orphan methyltransferases and those from restriction modification systems (RMSs) have been co-opted to regulate virulence epigenetically in many bacteria. However, the potential regulatory role of DNA methylation mediated by archetypal Type I systems in *Escherichia coli* has never been studied. We demonstrated that removal of DNA methylated mediated by three different *Escherichia coli* Type I RMSs in three distinct *E. coli* strains had no detectable effect on gene expression or growth in a screen of 1190 conditions. Additionally, deletion of the Type I RMS EcoUTI in UTI89, a prototypical cystitis strain of *E. coli*, which led to loss of methylation at >750 sites across the genome, had no detectable effect on virulence in a murine model of ascending urinary tract infection (UTI). Finally, introduction of two heterologous Type I RMSs into UTI89 also resulted in no detectable change in gene expression or growth phenotypes. These results stand in sharp contrast with many reports of RMSs regulating gene expression in other bacteria, leading us to propose the concept of “regulation avoidance” for these *E. coli* Type I RMSs. We hypothesize that regulation avoidance is a consequence of evolutionary adaptation of both the RMSs and the *E. coli* genome. Our results provide a clear and (currently) rare example of regulation avoidance for Type I RMSs in multiple strains of *E. coli*, further study of which may provide deeper insights into the evolution of gene regulation and horizontal gene transfer.

**Author summary:** DNA methylation is perhaps the most common epigenetic modification, and it is commonly associated with gene regulation (in nearly all organisms) and virulence (particularly well studied in bacteria). Regarding bacterial virulence, the current DNA methylation literature has focused primarily on orphan methyltransferases or phasevariable restriction modification systems (RMSs). Interestingly, no reports have studied the potential regulatory role of the first RMS discovered, the Type I RMS EcoKI. We used transcriptomics, Phenotype Microarrays, and a murine model of urinary tract infection to screen for functional consequences due to Type I methylation in three unrelated strains of *E. coli*. Remarkably, we found zero evidence for any epigenetic regulation mediated by these Type I RMSs. Thus, these Type I RMSs appear to function exclusively in host defense against incoming DNA (the canonical function of RMSs), while the methylation status of many hundreds of the corresponding recognition sites has no detectable impact on gene expression or any phenotypes. This led us to the concept of “regulation avoidance” by such DNA methyltransferases, which contrasts with the current literature on bacterial epigenetics. Our study hints at the existence of an entire class of regulation avoidant systems, which provides new perspectives on methylation-mediated gene regulation and bacterial genome evolution.

## Introduction

Regulation and response are fundamental aspects of cellular life. Epigenetics in the context of regulation refers to heritable changes in gene expression without alterations to the primary DNA sequence of an organism (1). This rapidly growing field encompasses an expanding list of biochemical incarnations spanning the entire central dogma (DNA and RNA modifications; chromatin remodelling; noncoding RNAs; and prion proteins) that adds additional layers of complexity and regulatory potential on top of primary DNA sequences (2–5). Epigenetics thus allows a clonal population to generate phenotypic heterogeneity, which can serve specific biological functions (6,7). Particularly for pathogens (and especially facultative pathogens), epigenetics is one way to implement rapid responses to new environments such as different host niches and immune pressures, which in turn may enhance survival and virulence (8,9).

DNA methylation represents the most extensively studied epigenetic mechanism in all domains of life (2,10–12). In eukaryotes, DNA methylation has been demonstrated to play an important role in differentiation, development and disease (including cancer) (13,14). Bacterial DNA methylation, in particular, has a special place in the collection of epigenetic implementations, as its role “outside genetics” in mediating phage resistance was discovered and characterized prior to the modern usage of the term epigenetics (15).

Bacterial DNA methylation occurs most commonly on adenine (N6-methyladenine, 6mA), but also on cytosine (C5-methylcytosine, 5mC and N4-methylcytosine, 4mC) nucleotides (16). The majority of known bacterial DNA methyltransferases belong to two general categories, restriction modification system (RMS)-associated and orphan methyltransferases (4). RMSs have been characterized as bacterial innate immune systems, using methylation of specific motifs to differentiate between self and non-self DNA (15). RMSs can be found in 90% of all sequenced prokaryotic genomes, with 80% possessing multiple systems, hinting at a diverse reservoir of potential epigenetic information (11). RMSs are classified based on co-factor requirement, subunit composition, cleavage pattern, and mechanism into 4 types (I to IV) (17–20). Type I RMSs were the first discovered and are multi-subunit enzyme complexes consisting of three proteins: a methyltransferase (typically denoted HsdM), an endonuclease (HsdR), and a sequence recognition protein (HsdS). Type I RMSs recognise bipartite sequences (for example, 5’-AAC(N_6_)GTGC-3’, where N=A,T,G or C) with each HsdS subunit possessing two highly variable target recognition domains (TRDs) which specify the two halves of the bipartite motif (19).

Beyond immunity, RMS-associated methyltransferases are also known to regulate gene expression. A dramatic example of such regulation is found in phasevariable Type I and Type III RMSs. Phase variation of these RMSs results in rapid and reversible changes to methylation patterns, resulting in coordinated changes to the expression of distinct sets of genes (regulons), hence the term “phasevariable regulons” or phasevarions (8,21). Phasevarions can be identified by known markers of phase variation, such as simple sequence repeats (SSRs) within promoters or gene bodies and *hsdS* genes flanked by inverted repeats. Systematic surveys have identified 13.8% Type I and 17.4% Type III RMSs as being potentially phase variable (22–24). Several clinically relevant pathogens such as *Helicobacter pylori* (25,26), *Neisseria meningitidis* (27), *Neisseria gonorrhoeae* (28,29), *Haemophilus influenzae* (30,31), *Moraxella catarrhalis (32,33),* and *Kingella kingae* (34) harbor such phase variable RMSs. An extreme example is found in *Streptococcus pneumoniae*, where some strains carry a phase variable Type I RMS which can result in up to six distinct methylation specificities, with different methylation patterns specifically associated with either invasive disease or nasopharyngeal carriage *in vivo (35–37).*

In contrast, orphan methyltransferases, as their name suggests, are methylating enzymes without a functional cognate endonuclease. Unlike RMSs, orphan methyltransferases are phylogenetically more conserved across genera/families (11,38,39) and even essential in certain bacteria (40–42). Orphan methyltransferases are not involved in host defense; instead, they play key housekeeping regulatory roles, ranging from DNA replication, mismatch repair, controlling transposition, and gene expression (43). Regarding gene expression, mutation of these orphan methyltransferases can have large effects on the transcriptome, affecting hundreds to thousands of genes (44–49); not surprisingly, therefore, mutation of orphan methyltransferase can also lead to defects in virulence (50–55). In some cases, there exist locus-specific roles of methylation in virulence factor regulation, such as for P pili and the autotransporter antigen 43 (Agn43) (56,57). The two most well-studied orphan methyltransferases are Dam (DNA adenine methyltransferase) in gamma-proteobacteria and CcrM (cell cycle regulated DNA methyltransferase) in alpha-proteobacteria (58).

Therefore, DNA methylation can generally affect both gene expression and phenotypes, of which virulence is an especially interesting case in pathogens. Interestingly, there has been a relative dearth of such studies on the archetypal Type I RMSs, which are typically non-phase variable and retain both restriction and modification functions. We used single molecule sequencing to characterize the Type I RMS in a uropathogenic *Escherichia coli* strain, UTI89. We verified that this Type I RMS was functional for restriction of incoming transformed DNA. Surprisingly, we could find no effect on gene expression or virulence when this RMS was removed, despite >700 DNA bases changing in methylation state. Similar results were found when we removed the Type I RMS from two other *E. coli* strains, MG1655 and CFT073. Intrigued by these finding, we replaced the native UTI89 Type I RMS with the Type I RMS from MG1655 and CFT073; in both cases, re-installing methylation at ~400-600 distinct sites in the genome also had no effect on gene expression or on a broad panel of bacterial growth phenotypes. These data suggest that, in *E. coli,* these canonical Type I systems are purely host defense mechanisms devoid of any secondary regulatory functions in host physiology and virulence. This is the first reported example of “regulation avoidance” for any DNA methylation system in bacteria. The fact that three distinct methylation specificities all demonstrate this regulation avoidance raises further speculations about the evolution and plasticity of both the genome and the Type I RMSs in *E. coli*.

## Results

### Uropathogenic *Escherichia coli* UTI89 possesses a functional Type I Restriction Modification System

The clinical cystitis strain UTI89 encodes a Type I RMS at the *hsdSMR* locus (59,60). PacBio single molecule real time (SMRT) sequencing identified 3 methylation motifs in UTI89: 5’-GATC-3’, 5’-CCWGG-3’, and 5’-CCA(N_7_)CTTC-3’ (S1 Table). The first two motifs are well known in *E. coli* as the methylation sites of orphan methyltransferases Dam and Dcm, respectively (12,61). The third motif, a bipartite N6-methyladenine (m6A) motif, is typical for a Type I RMS (19). Deletion of the *hsdSMR* locus in UTI89 resulted in loss of adenine methylation only at the 5’-CCA(N_7_)CTTC-3’ motif, confirming the specificity of this RMS (S1 Table).

To verify that the UTI89 *hsdSMR* locus was functional for restriction of incoming DNA, we performed a classic plasmid transformation efficiency assay (62). A plasmid containing the 5’-CCA(N_7_)CTTC-3’ motif was isolated from the UTI89*ΔhsdSMR* strain (which does not methylate this motif); this plasmid was transformed less efficiently into wild type (wt) UTI89 than into an otherwise isogenic UTI89*ΔhsdSMR* strain (Fig 1A). Moreover, as seen with other Type I RMSs, addition of a second motif further reduced the efficiency of transformation into wt UTI89 (Type I RMSs cleave DNA when translocation is stalled due to collision between adjacent Type I complexes (63)). When plasmids were isolated from wt UTI89 (which methylates the 5’-CCA(N_7_)CTTC-3’ motif), no difference in efficiency was seen between transformations into UTI89 and UTI89*ΔhsdSMR* (Fig 1A). Thus, uropathogenic *E. coli* UTI89 possesses a functional Type I RMS, with a formal name of RM.EcoUTI and a specificity of 5’-CCA(N_7_)CTTC-3’ (64). We hereafter refer to this system as EcoUTI.

**Fig 1.**
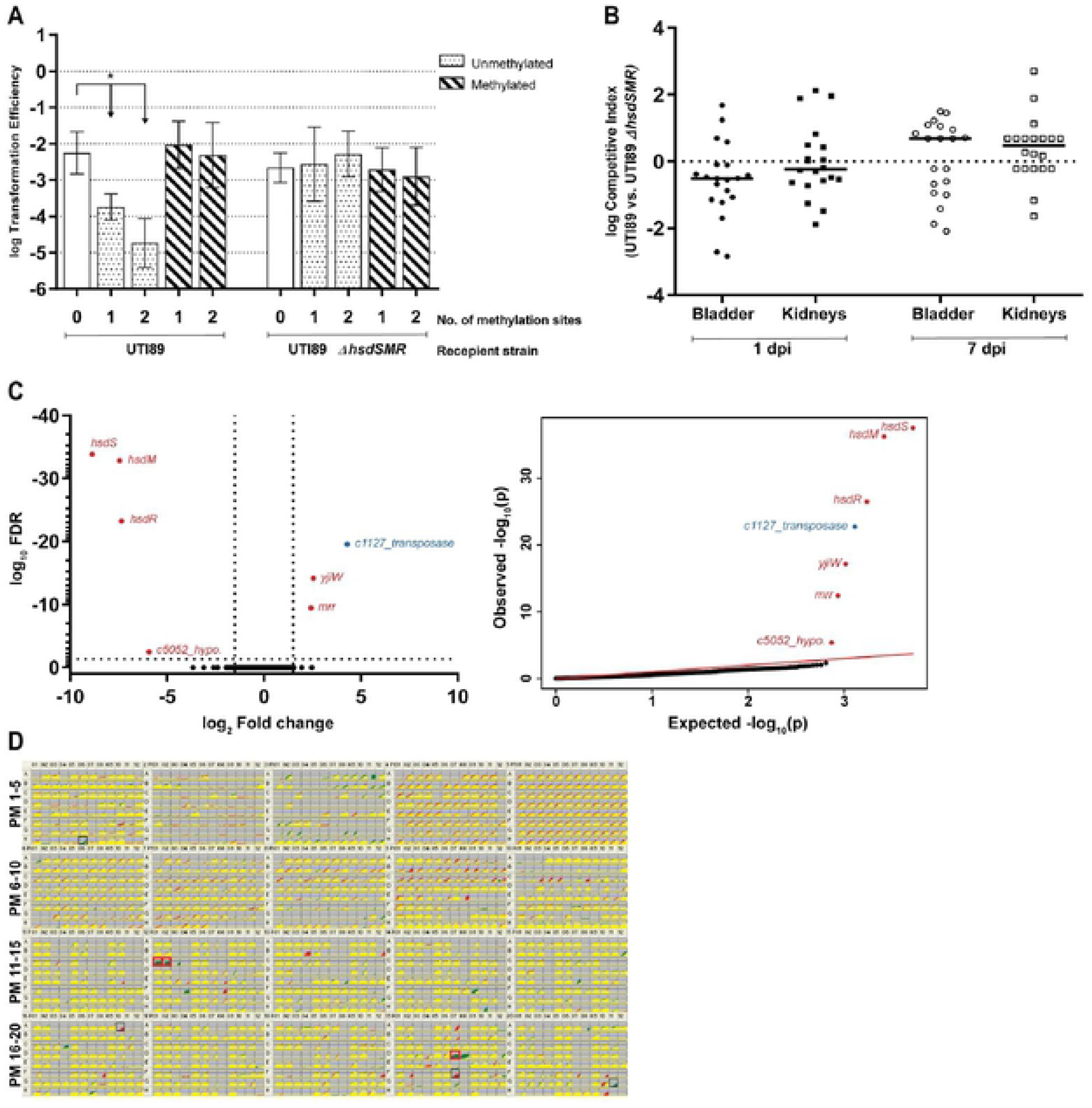
UTI89 Type I RMS EcoUTI functions only as a restriction system, with no apparent regulatory role. **(A)** Transformation efficiency assay using plasmids bearing 0, 1, or 2 copies (as indicated on the x-axis) of the UTI89 Type I RMS motif (5’-CCA(N_7_)CTTC-3’). Recipient cells were wild type UTI89 and the isogenic *ΔhsdSMR* mutant, as indicated by the labels below the x-axis. Unmethylated and methylated plasmid preparations were used to transform each strain, as indicated by the legend at the top right. An unpaired t-test was used to identify significant differences between plasmids with 0, 1 and 2 methylation sites for both preparations and strains; * p <0.05, n = 3 biological replicates. Data represents mean ± standard deviation (s.d.) of log transformed values. **(B)** Competitive index (CI) for in vivo co-infections with wt UTI89 and UTI89*ΔhsdSMR*. Bladder and kidney pairs (as indicated on the x-axis) were aseptically harvested at 1 or 7 days post infection (dpi), as indicated by the labels below the x-axis, and plated on appropriate selective plates for calculation of CI. A Wilcoxon signed rank test was used to test for a significant difference of log CI from 0; * p <0.05, n = 20 mice/time point, performed as 2 biological replicates. Each point represents data from a single mouse; horizontal lines represent the median. **(C)** RNA sequencing comparing logarithmic phase transcriptomes of UTI89 and UTI89*ΔhsdSMR.* Left, a volcano plot of log FDR against log fold change. Right, qq-plots showing the distribution of uncorrected p-values. Significantly differentially expressed genes (log_2_ fold change >1.5 and log_10_ FDR <0.05) are labelled and colored either red (deleted genes/polar effects) or blue (validated as false positive by qRT-PCR). n = 3 biological replicates. **(D)** Phenotype microarray (PM) panel with plates PM1 to 20 comparing wt UTI89 and UTI89*ΔhsdSMR*. Each plate is represented as a 12×8 grid of growth curves (red (wt), green (*ΔhsdSMR*), and yellow (overlap) on a gray background). Each growth curve plots growth (measured colorimetrically) (y-axis) against time (x-axis). Wells representing conditions where a height difference was observed between the strains in both replicates are boxed in black, and wells which also have a quality score >150 are considered significant and boxed in red. n = 2 biological replicates.

### EcoUTI Type I methylation has no detectable role during *in vivo* urinary tract infection

DNA methylation is widely recognized to influence gene expression and virulence in multiple bacterial pathogens (51,58,65). EcoUTI methylates 754 sites in the UTI89 genome; we therefore hypothesized that complete removal of this system would affect both gene expression and virulence. The UTI89 and UTI89*ΔhsdSMR* strains had no difference in growth in both rich and minimal media nor in Type I pilus expression *in vitro* (data not shown). We then tested virulence using a transurethral murine model for ascending urinary tract infection (UTI) (66). Competitive co-infections in 6-8 week old C3H/HeN mice using equal mixtures of wild type UTI89 and UTI89*ΔhsdSMR* showed no competitive advantage for either strain at 1 or 7 days post-infection (dpi) in either bladders or kidneys (Fig 1B). Comparison of bacterial loads from single infections also showed no significant difference between the two strains (S1 Fig).

### EcoUTI Type I methylation has no effect on gene expression or growth in multiple conditions

The lack of an *in vivo* infection phenotype, despite the complete removal of EcoUTI methylation, contrasts with reports on multiple other methylation systems (including RMSs) that impact bacterial phenotypes (47,67–71). We therefore asked whether changing methylation could alter any gene expression using RNA-seq. When comparing UTI89 with the otherwise isogenic UTI89*ΔhsdSMR* strain, in both log and stationary phase in rich media, only 7 genes were significantly changed in expression in the mutant: the (deleted) *hsdSMR* and UTI89_C5052 genes were downregulated, the two genes flanking the *hsdSMR* locus were upregulated, and a single transposase gene was upregulated. In the UTI89 genome, UTI89_C5052 is a hypothetical gene annotated in the intergenic region between *hsdM* and *hsdR*, so it was deleted with the entire *hsdSMR* locus. Altered expression of the two flanking genes was connected to the use of an antibiotic resistance cassette to knock out *hsdSMR* (see methods). Targeted qRT-PCR on the transposase gene was unable to validate the change seen in RNA-seq (Fig 1C, S2 Fig, S2 and S3 Tables). Therefore, we could find no change in gene expression that was attributable to loss of Type I methylation.

Although RNA-seq is a powerful tool for assaying gene expression, it only tests expression in one experimental growth condition. To test a broader range of growth conditions, we used Biolog Phenotype Microarrays (PM) (72). PM plates 1-20 represent 1190 different nutrients, toxins, antibiotics, inhibitors, and other conditions. Relative to wt UTI89, the UTI89*ΔhsdSMR* strain had gained resistance to the aminoglycoside antibiotic paromomycin (PM12, wells C01-02), which was consistent with the use of a kanamycin resistance cassette to knock out the *hsdSMR* locus. (Fig 1D, S4 Table). UTI89*ΔhsdSMR* had also gained resistance to the formazan dye Iodonitrotetrazolium (INT) violet (PM19, well D07). There are four wells containing different concentrations of INT on PM19 (D05 - D08); well D07 is the second-lowest concentration, and the other three wells were not called as different from wt UTI89. In particular, at the two highest concentrations (wells D05 and D06) there was very little difference in the growth curves between UTI89 and UTI89*ΔhsdSMR.* We therefore suspect that this was a false positive phenotype (Fig 1D, S4 Table). We thus find, strikingly, no phenotypic difference attributable to the loss of methylation.

### Loss of Type I methylation in two other *E. coli* strains also has no effect on gene expression or growth phenotypes

We next asked whether the lack of any detectable gene expression or growth phenotype changes was unique to UTI89 and the EcoUTI methylation system. *E. coli* strain MG1655 is a well-studied, lab-adapted K12 strain, while CFT073 is a commonly used pyelonephritis strain; both also carry a single Type I RMS (EcoKI and EcoCFTI, respectively). EcoKI methylates the 5’-AAC(N_6_)GTGC-3’ motif, while EcoCFTI is predicted to methylate the 5’-GAG(N_7_)GTCA-3’ motif (59,73). Plasmid transformation efficiency assays confirmed that both EcoKI and EcoCFTI also were functional as restriction systems for incoming DNA, targeting the previously reported motifs (S3 Fig). In RNA-seq experiments, besides the expected change in expression of the *hsdSMR* genes themselves and the flanking genes affected by insertion of the antibiotic resistance cassette, MG1655*ΔhsdSMR* had 4 genes upregulated during log phase, all encoded within a predicted cryptic prophage and validated by qRT-PCR (Fig 2A, S4A Fig, S2 and S3 Tables). However, independently generated clones of the MG1655*ΔhsdSMR* strain did not have a similar upregulation of these four prophage genes when tested by qRT-PCR; we therefore consider the expression changes in these four genes an artefact of that single strain, possibly due to a second site mutation during generation of the knockout (data not shown). The CFT073*ΔhsdSMR* strain had limited changes in gene expression relative to the wild type CFT073, including the *hsdSMR* genes, and flanking gene *yjiW* (polar effect). Targeted qRT-PCR on *malK* and *lamB* genes was unable to validate the changes seen in RNA-seq, confirming these as false positives (Fig 2B, S4B Fig, S2 and S3 Tables).

**Fig 2.**
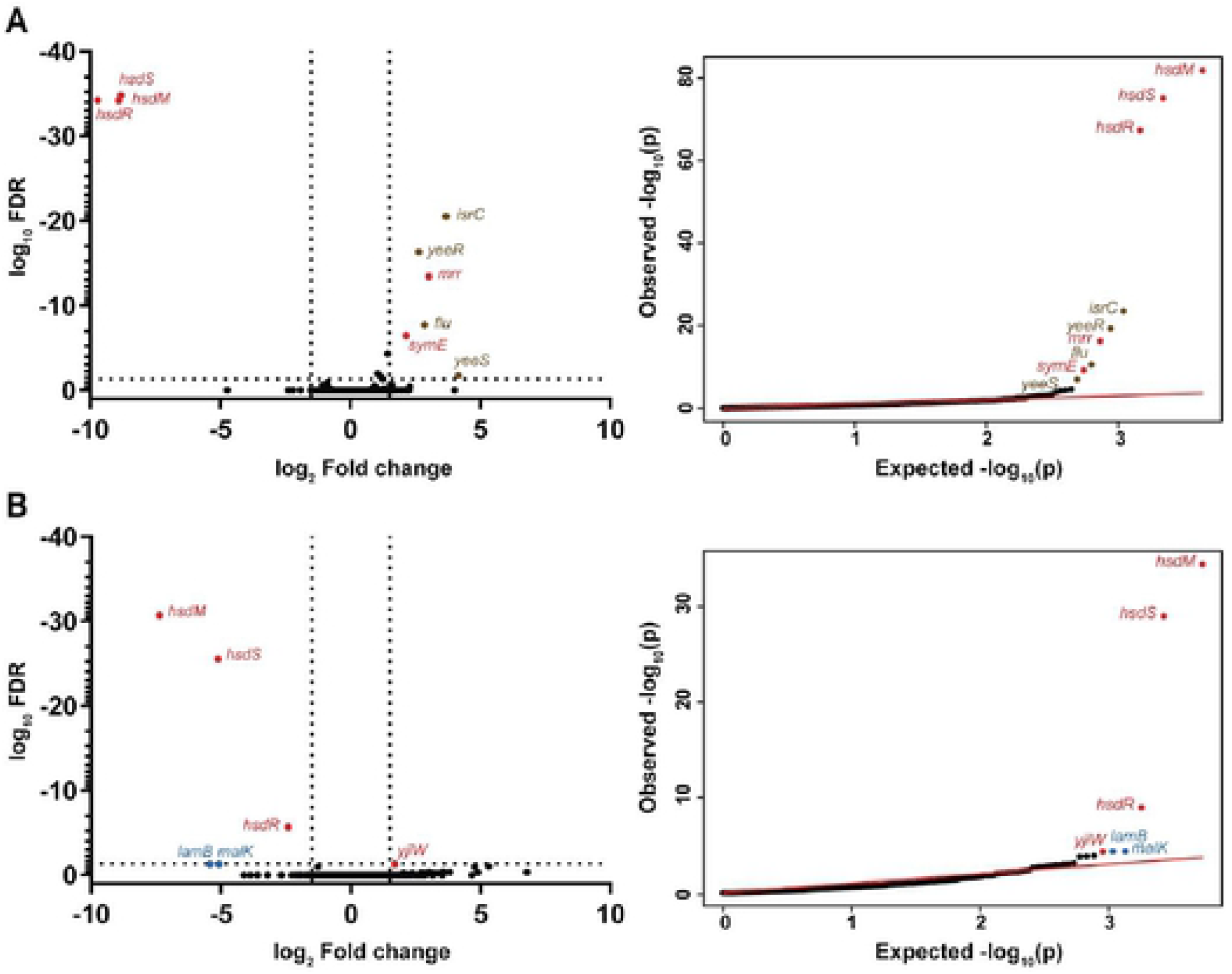
Type I RMSs from *E. coli* CFT073 and MG1655 do not affect gene expression. RNA sequencing comparing logarithmic phase transcriptomes of **(A)** MG1655 and MG1655*ΔhsdSMR*, and **(B)** CFT073 and CFT073*ΔhsdSMR.* Left, a volcano plot of log FDR against log fold change. Right, qq-plots showing the distribution of uncorrected p-values. Significantly differentially expressed genes (log_2_ fold change >1.5 and log_10_ FDR <0.05) are labelled and colored either red (deleted genes/polar effects), brown (differentially expressed genes) or blue (validated as false positive by qRT-PCR). n = 2 (**A** MG1655) or 3 (**B** CFT073) biological replicates.

Using the Biolog PM, we again found no phenotypic differences between either (i) MG1655 and MG1655*ΔhsdSMR* or (ii) CFT073 and CFT073*ΔhsdSMR*; except for gained antibiotic resistance due to kanamycin resistance cassette used to knock out the *hsdSMR* locus (S4 Table, S4C and S4D Figs). Thus, in three distinct *E. coli* strains, each encoding Type I RMSs with different specificities, deletion of the entire RMS had no detectable effect on gene expression or any growth phenotypes we measured.

### Switching Type I methylation systems in UTI89 also does not affect gene expression or any growth phenotypes

Type I RMSs can be highly polymorphic among different strains of the same species. In many cases, this is thought to be due to both mutation and recombination, possibly driven by diversifying selection (74,75). Furthermore, the whole *hsdSMR* locus need not be recombined; Type I RMSs are classified into 5 families (designated A through E) based on the similarity of the *hsdM* and *hsdR* genes, enabling intra-family genetic complementation with divergent *hsdS* genes (19). As *hsdS* encodes the specificity determinant, mutation or recombination of just the *hsdS* gene is sufficient to alter the methylation and restriction specificity of the entire system (35,76). The RMSs in UTI89 and MG1655 are both subclassified as Type IA and demonstrate this latter relationship; the *hsdM* and *hsdR* genes in these two strains have very similar sequences (99% identical), while the *hsdS* genes are only 45.4% identical and direct distinct specificities (Fig 3A). The Type I RMS in CFT073, on the other hand, is a Type IB RMS with <40% identity to the Type IA system in all three genes (Fig 3A).

**Fig 3.**
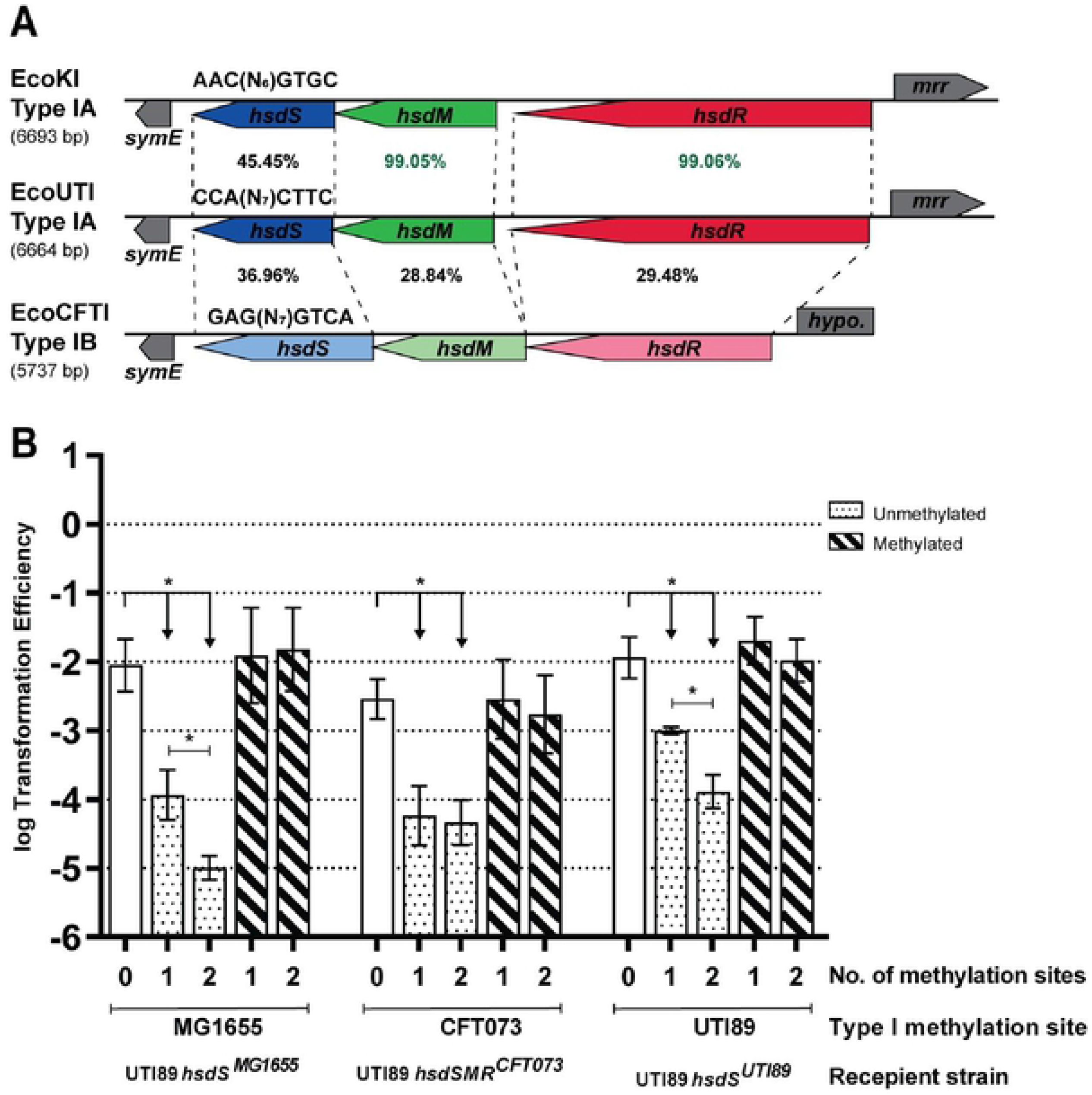
Installation of different Type I methylation specificities into UTI89. **(A)** Graphical representation of the host specificity determinant (*hsd*) locus showing the amino acid identity between Type I RMSs EcoKI, EcoUTI, and EcoCFTI from *E. coli* strains MG1655, UTI89, and CFT073 respectively. The subclassification for each system is indicated on the left, with the total length of DNA spanned by the *hsd* genes shown in parentheses. The *hsd* genes are shown in different colors for visualization and drawn to scale, with transcription direction indicated by the arrow. Dotted lines further connect the corresponding beginnings and ends of each *hsd* gene in different systems. Genes adjacent to the *hsd* genes are shown in gray. The percentage identity between *hsd* genes from different systems is indicated; green >99% and black <50%. **(B)** Transformation efficiency assay using plasmids bearing 0, 1, or 2 copies (as indicated on the x-axis) of the MG1655 (5’-AAC(N_6_)GTGC-3’), CFT073 (5’-GAG(N_7_)GTCA-3’) and UTI89 (5’-CCA(N_7_)CTTC-3’) Type I RMS motifs. Recipient cells were UTI89 *hsdS^MG1655^*, UTI89 *hsdSMR^CFT073^* and UTI89 *hsdS^UTI89^*, as indicated by the labels below the x-axis. Unmethylated and methylated plasmid preparations were used to transform each strain, as indicated by the legend at the top right. An unpaired t-test was used to identify significant differences between plasmids with 0, 1 and 2 methylation sites for both preparations and strains; * p <0.05, n = 3 biological replicates. Data represents mean ± s.d. of log transformed values.

To analyze the effect of changing Type I methylation specificity, we created two derivatives of UTI89: one where the *hsdS* gene was replaced by the MG1655 *hsdS* allele (UTI89 *hsdS^MG1655^*), and one where the entire *hsdSMR* locus was replaced by the EcoCFTI locus (UTI89 *hsdSMR^CFT073^*). As a control, we also re-inserted the UTI89 *hsdS* allele (UTI89 *hsdS^UTI89^*) using the same cloning strategy as that used to make UTI89 *hsdS^MG1655^* (i.e. UTI89 *hsdS^MG1655^* and UTI89 *hsdS^UTI89^* share the same parental strains and have undergone the same cloning steps). PacBio SMRT sequencing confirmed that plasmids isolated from each of these strains methylated only the expected motif (data not shown). Plasmid transformation efficiency assays also showed that all 3 “methylation-switch” strains generated indeed had functional Type I RMSs with specificities as expected based on the encoded *hsdS* allele (i.e. UTI89 *hsdS^MG1655^*, UTI89 *hsdSMR^CFT073^*, and UTI89 *hsdS^UTI89^* restricted only plasmids carrying the EcoKI, EcoCFTI, and EcoUTI motif respectively, in a methylation-dependent manner) (Fig 3B).

These strains introduce two different Type I methylation specificities in the UTI89 genome, accounting for 641 (MG1655) and 423 (CFT073) predicted newly methylated sites (Fig 4A, S5 Table). All together, the three RMSs we examined account for 1818 distinct methylation sites that would vary in methylation status among the strains we studied. The number of intergenic methylation sites in UTI89, which are expected to be more likely to affect gene regulation (46,77–80), are comparable for the three Type I RMSs (31-40 intergenic sites) (S5 Table). Using RNA-seq, we again found no gene expression changes for any of the “methylation-switch” strains compared with the parental wt UTI89, except for one hypothetical gene that was close to both the FDR and fold-change cutoffs (Fig 4C and 4D, S5 Fig, S2 and S3 Tables). Furthermore, Biolog PM also showed no significant changes attributable to Type I methylation in UTI89 *hsdS^MG1655^* (S4 Table, S6 Fig). Notably, there was no difference in INT resistance between UTI89 *hsdS^MG1655^* and wt UTI89 at any concentration, consistent with our previous interpretation that the single difference identified between UTI89 and UTI89*ΔhsdSMR* was a false positive (Fig 1D, S4 Table).

**Fig 4.**
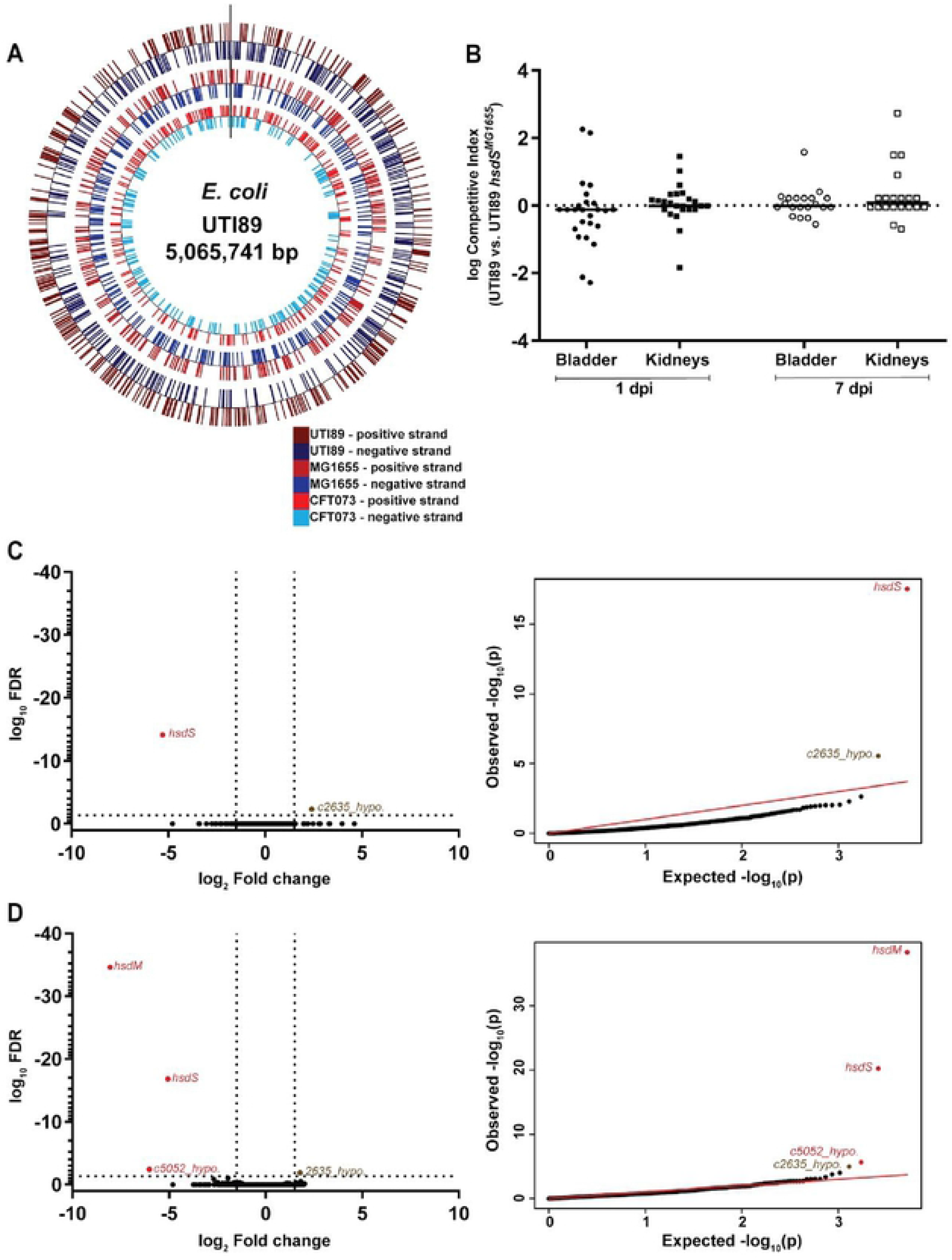
Heterologous Type I methylation in UTI89 does not affect virulence and gene expression. **(A)** Distribution of heterologous Type I methylation sites in the UTI89 genome. Methylation sites for different systems are indicated by tick marks on different concentric circles; from the outer to the inner circle, UTI89 (747 sites), MG1655 (628 sites), and CFT073 (415 sites) sites are represented. Positions on the positive and negative strands are represented by colored tick marks as indicated in the legend at the bottom right, marked in red and blue respectively. **(B)** Competitive index (CI) for in vivo co-infections with wt UTI89 and UTI89 *hsdS^MG1655^*. Bladder and kidney pairs (as indicated on the x-axis) were aseptically harvested at 1 or 7 days post infection (dpi), as indicated by the labels below the x-axis, and plated on appropriate selective plates for calculation of CI. A Wilcoxon signed rank test was used to test for a significant difference of log CI from 0; * p <0.05, n = 25 mice (1dpi) and 20 mice (7dpi), performed as 2 biological replicates. Each point represents data from a single mouse; horizontal lines represent the median. **(C and D)** RNA sequencing comparing logarithmic phase transcriptomes of **C)** UTI89 and UTI89 *hsdS^MG1655^*, and **D)** UTI89 and UTI89 *hsdSMR^CFT073^.* Left, a volcano plot of log FDR against log fold change. Right, qq-plots showing the distribution of uncorrected p-values. Significantly differentially expressed genes (log_2_ fold change >1.5 and log_10_ FDR <0.05) are labelled and colored either red (deleted genes/polar effects) or brown (differentially expressed genes). n = 3 biological replicates.

As a final functional test, we used several *in vitro* and *in vivo* assays for virulence. Changing the methylation system had no effect on growth rate in rich or minimal media or on virulence-related assays such as motility and biofilm formation (S7 - S9 Figs). Competitive co-infections with UTI89 and UTI89 *hsdS^MG1655^* again revealed no competitive advantage for either strain, irrespective of the time-point or organ tested (Fig 4B).

To test whether UTI89 was unique in its ability to tolerate changes in methylation, we created a similar “methylation-switch” in MG1655 by inserting the UTI89 *hsdS* allele. We again found no differences between wt MG1655 and MG1655 *hsdS^UTI89^* in the Biolog PM phenotype screen (S4 Table, S10 Fig). We therefore conclude that, at least in UTI89 and MG1655, substantial changes in DNA methylation across the genome are not merely tolerated but simply have no measurable effect on gene expression or laboratory-measured phenotypes.

## Discussion

The EcoKI restriction modification system (RMS) was identified over a half century ago due to its role in the specific restriction of exogenous (phage) DNA (73). EcoKI is thus the prototypical example of a RMS, but subsequent studies also demonstrated that, particularly for *E. coli* but also for many other bacteria, this was an archetypal system as well. It was thus designated “Type I” in the RMS classification that followed. As more RMSs were discovered, they were initially assumed to play similar roles in host defense and horizontal gene transfer (81,82). In bacteria, one of the early indications that methylation could affect transcription arose from the study of regulation of RMSs themselves, some of which utilize methylation as a readout of expression level, effectively a type of product-inhibition feedback (83). The general observation that DNA methylation, particularly that mediated by orphan methyltransferases (such as Dam or CcrM, which may themselves have evolved from RMSs), could alter transcription (44,48,84), led to the discovery of a broad suite of associated roles in DNA metabolism, cell physiology, and virulence (12,51,55,85–87). An active research community continues to describe and characterize the additional roles that DNA methylation (from both restriction and orphan systems) play in bacteria (4,39,88), some of which can be very specific (67,69). We now show that, in contrast with expectations from this literature, the archetypal EcoKI Type I RMS has nearly zero impact on gene regulation or any phenotype among more than 1000 growth conditions tested. These results generalize to two additional Type I RMSs from pathogenic strains of *E. coli*. We therefore conclude that these Type I RMSs have a strictly limited role in the classical defense against incoming foreign DNA, and the associated methylation of hundreds of adenines throughout the genome specifically avoids any consequential regulation of gene expression.

The paradigm of “one gene, one function” made the initial discovery that restriction systems could play a distinct epigenetic role in gene regulation somewhat surprising. Now, the idea that methylation can affect the interaction with DNA binding proteins is quite clear (12), and orphan methyltransferases were thought to have preserved DNA methylation for such non-defense (i.e. regulatory) functions (11). Perhaps the strongest evidence that gene regulation is a biologically selected function for RMSs comes from the discovery of phasevarions, in which phase variable expression (through recombination or simple sequence repeat variation) leads to rapid switching between RMSs with different methylation specificities, leading to differential regulation of genes that have particularly been shown to impact virulence (8). Type I systems are the least common of the four classes of RMSs, however, and generally have the longest recognition sequences (leading to fewer modified sites in a given genome) (11,39,59). Regardless, phasevarions composed of multiple Type I specificities have also been described, with similar effects on gene expression and virulence (35,36,89); in addition, non-phase variable Type I systems have also been shown to affect gene expression (67–69).

The three Type I RMSs studied here, EcoUTI, EcoCFTI, and EcoKI, each methylate many hundreds (427 to 754) of distinct sites in their respective host genomes (and similar predicted numbers in other *E. coli* genomes). We tested for phenotypic impacts using an *in vivo* murine model of urinary tract infection, a complex phenotype which demands that inoculated cells react to changing nutrient, fluid flow, and host immune conditions. In addition, *E. coli* are known to occupy different niches within the urinary tract, ranging from bladder epithelial cells (both within the cytoplasm and on the surface) to the lumen of the bladder and the parenchyma of the kidney (90). While we did not investigate these different niches individually, there was no overall difference in infection titers, even during competition experiments with a wild type strain, a strong indication that no virulence defect is present when EcoUTI is deleted. Complementing this complex phenotype, we used the Biolog PM platform to screen a “simple” growth phenotype in 1190 conditions, again finding no difference regardless of the presence or absence of the three Type I RMSs. Finally, we used RNA-seq to explore whether gene expression (either on a global scale or at the level of individual genes) was altered by changing methylation, again finding no differences attributable to DNA methylation. It remains possible that under some specific conditions, these Type I systems would indeed alter gene expression or phenotypes; indeed, methyltransferases such as *Escherichia coli* Dcm and *Mycobacterium tuberculosis* MamA alter gene expression primarily under specific growth conditions such as stationary phase and hypoxia, respectively (61,78,91). However, it seems that finding such a condition might serve as the “exception that proves the rule”, given our data using both simple and complex phenotypes, across >1000 growth conditions, and measuring genome-wide gene expression under lab conditions. Notably, in the course of making one of the Type I RMS knockouts, we recovered a clone carrying a nonsynonymous mutation in a nucleotide biosynthesis operon; control experiments using this clone demonstrated that we could indeed detect differences in *in vivo* infections, PM arrays, and gene expression (manuscript in preparation).

Based on our results, we propose the concept of “regulation avoidance”. This term describes a state where methylation (i.e. changing from unmethylated to methylated or vice versa) has no measurable impact on gene expression or any phenotype. We have shown that the Type I RMSs in UTI89, MG1655, and CFT073 all possess the property of regulation avoidance in their respective strains; furthermore, the MG1655 EcoKI and CFT073 EcoCFTI RMSs have regulation avoidance when introduced into UTI89, and the UTI89 EcoUTI RMS has regulation avoidance when introduced into MG1655. We strongly suspect that regulation avoidance is not limited to the specific *E. coli* strains and Type I systems studied here, despite the fact that many RMSs have been shown to not have this property (65,67–69). Therefore, future studies will be required to characterize the regulatory role of other RMSs, potentially leading to general principles that may help explain when and why some RMSs have regulation avoidance (possibly only in some strains). At this point, we suspect that Type I systems, due to their longer recognition sequence and overall fewer methylated sites, will be more likely to be regulation avoidant than other RMSs.

A few additional implications of regulation avoidance merit discussion here. First, regulation avoidance likely is related to the fact that RMSs are mobile, or that they switch regularly, on an evolutionary time scale. A permanent and fixed pattern of methylation would likely result in adaptation of the host genome and host proteins to accommodate those sites where methylation overlaps binding sites for transcription factors or RNA polymerase. This would be similar to the process of amelioration of codon usage or compensatory mutations to minimize fitness defects from new plasmids, for example (92,93). Our data, showing that removal and replacement of each of three separate Type I systems has no detectable impact on regulation in UTI89, instead implies that any genomic adaptation would have occurred towards *both* methylated and unmethylated motifs. Second, some *E. coli* strains indeed do not have a Type I RMS, but the majority do; furthermore, RMSs are thought to be easily transferred between strains (also on an evolutionary time scale) (74,75). In particular, details of the structure of Type I RMS enable rapid switching of specificity, for example by evolution of or limited recombination of the target recognition domain of the *hsdS* gene (19,65). Our data on strains in which we installed two different Type I RMSs into UTI89 is consistent with this notion, and we speculate that the lack of any effect in deleting EcoUTI is actually driven by the presence of other Type I RMSs in the evolutionary history of UTI89. specificity through mutation of the *hsdS* gene, regulation avoidance is possibly also mediated by adaptation of the recognition sequence of Type I RMSs. Finally, early examples of methylation regulating gene expression in bacteria came from studies of the regulation of expression of the RMSs themselves. One consistent theme from such studies is that some details of the regulation can be rationalized using a scenario where a new RMS is introduced into a bacterium: methylation is expressed first, while restriction is expressed only with a delay, thus preventing immediate degradation of the chromosome (94). This is similar to the requirements expected for introduction of a new toxin-antitoxin system (requiring antitoxin expression before toxin expression) (95), and further suggests that the introduction of a new RMS may be a regular occurrence. Of note, we have emphasized the evolutionary time scale here to contrast with phasevarions, which switch RMS specificity on a time scale much shorter than that required for genomic adaptation.

In summary, we have conclusively demonstrated that, for 3 different strains of *E. coli*, the Type I RMSs have very nearly zero effect on gene expression and no effect on any phenotype tested besides their originally described function of defense against incoming foreign DNA. We introduce the concept of “regulation avoidance” to highlight the contrast of our results with the weight of the published literature on non-defense functions of other RMSs and orphan methyltransferases. Regulation avoidance, in turn, leads to several insights into the evolutionary dynamics of RMSs, which suggest fruitful avenues for future study. In particular, how general is regulation avoidance among *E. coli*, and is it limited to Type I RMSs? Do other bacteria have regulation avoidant RMSs? Is regulation avoidance necessarily dependent upon preservation of DNA restriction function, or is evolutionary plasticity and exchange of methylation specificity sufficient? How much is regulation avoidance mediated by adaptation of DNA binding proteins, placement and/or context of binding sites in the genome, or modification of RMS specificity? Future studies addressing these questions may provide further insights into the impact that RMSs (and, by implication, phages and other mechanisms that introduce foreign DNA) have had on bacterial genome evolution.

## Material and methods

### Ethics statement

All animal experiments were approved by and performed in strict accordance with the Institutional Animal Care and Use Committee (IACUC) of A*STAR (Protocols #110605 and #130853).

### Media and culture conditions

All strains were propagated in Lysogeny Broth (LB) at 37°C unless otherwise noted. Media was supplemented with ampicillin (100 μg*/*ml), kanamycin (50 μg*/*ml), or chloramphenicol (20 μg*/*ml) where required (all antibiotics from Sigma, Singapore). M9 minimal medium (1X M9 salts, 2mM Magnesium sulphate, 0.1mM Calcium chloride, 0.2% glucose) and Yeast extract-Casamino acids (YESCA) medium (10g/L Casamino acids, 1g/L Yeast extract) were used for specific experiments.

### Strain and plasmid generation

Deletion mutants were generated using a λ-Red recombinase mediated strategy optimised for clinical strains of *E. coli*, with minor modifications (96). Briefly, a positive selection cassette encoding either kanamycin (*neo*) or chloramphenicol (*cat*) resistance was amplified from plasmid pKD4 or pKD3, respectively (97). Primers incorporated, at their 5’-end, 50bp of homology to the genomic locus being knocked out. The resultant PCR product was transformed into cells expressing λ-Red recombinase from vector pKM208, recovered at 37°C for 2 hours with shaking followed by 2 hours without shaking, then plated onto the appropriate antibiotic at 37°C overnight.

Strains containing a seamless marker-free replacement of a gene with a different allele were generated using a previously described negative selection strategy (98). Two successive rounds of λ-red recombinase mediated homologous recombination were performed, using the protocol described above, to first replace the wild type allele with a dual positive-negative selection cassette (amplified from plasmids pSLC217 or pSLC246 instead of pKD4 or pKD3). A second round of recombination was then performed using a PCR product with the desired allele instead of a resistance cassette, and the second selection done on M9 supplemented with 0.2% rhamnose. Correct clones were confirmed by PCR and by sequencing the recombination junctions by Sanger sequencing (1^st^ Base, Singapore).

The Type I RMS recognition motifs for UTI89 (5’-GAAG(N_7_)TGG-3’), MG1655 (5’-AAC(N_6_)GTGC-3’), and CFT073 (5’-GAG(N_7_)GTCA-3’) were incorporated into primers designed using an inverse PCR strategy to amplify plasmid pACYC184 (which does not natively possess any of these Type I recognition motifs). Linear plasmid amplicons were then digested using NotI, recircularized with T4 DNA ligase, then transformed into *E. coli* TOP10. When two copies of the Type I motifs were inserted, they were placed 1500bp apart (pACYC184 coordinates 381 and 1974). All strains, plasmids and primers (Sigma, Singapore) used in this study are listed in S6 and S7 Tables, respectively.

### Transformation efficiency assay

Competent cells were prepared using log phase cultures obtained by sub-culturing overnight bacterial culture 1:100 in LB and growing at 37°C up to OD_600_ = 0.4 – 0.5. Cells were washed twice with sterile water followed by a final wash with sterile 10% glycerol and resuspension in 1/100 of the original culture volume of 10% glycerol; these were then stored at −80°C in 50 μl aliquots. Plasmids with 1 or 2 copies of the bipartite Type I motif were extracted from the appropriate wild type or Type I RMS mutants to obtain methylated or unmethylated preparations. Plasmids were extracted using the Hybrid-Q plasmid miniprep kit (GeneAll, South Korea) according to the manufacturer’s recommended protocol. Competent cells were transformed with 100ng of each plasmid using 1mm electroporation cuvettes in a GenePulser XCELL system, at 400 Ω resistance, 25μF capacitance, and 1700V output voltage (Bio-Rad, Singapore). Cells were recovered in 1 ml of prewarmed LB at 37°C for 1 hour with shaking and plated on selective (chloramphenicol) and non-selective (LB) plates. Transformation efficiency was calculated by dividing the cfu/ml obtained on selective by that on non-selective plates per unit amount of plasmid DNA.

### Mouse infections

*In vivo* infections were performed using a murine transurethral model of urinary tract infection (66). Briefly, bacterial strains were grown in Type I pili-inducing conditions by two passages in LB broth at 37°C for 24 hours without shaking; a 1:1000 dilution was made from the first to the second passage. Cells were then harvested by centrifugation and resuspended in sterile cold PBS to OD_600_ = 1. Type I piliation for each strain was evaluated by a hemagglutination assay and Type I phase assay as described previously (99). A 1:2 dilution of the PBS suspension (final OD_600_ = 0.5) was then used as the inoculum. 7-8 week old female C3H/HeN mice (InVivos, Singapore) were anaesthetized using isoflurane and 50 μl of inoculum (OD_600_ = 0.5, ~1-2 × 10^7^ cfu/50 μl) was transurethrally instilled into the bladder using a syringe fitted with a 30 gauge needle covered with a polyethylene catheter (Product #427401, Thermo fisher scientific, USA). At specified times, mice were sacrificed and bladders and kidneys were harvested aseptically and homogenized in 1 ml and 0.8 ml of sterile PBS, respectively. Ten-fold serial dilutions were plated on appropriate selective plates to quantify bacterial loads. For co-infections, the inoculum consisted of a 1:1 mixture of two strains with an antibiotic resistance cassette inserted at the phage HK022 attachment site *attP* (100), each at 1-2 × 10^7^ CFU/50 μl; otherwise, the procedure was identical to that described for the single infections above. To account for potential bias due to selection markers, co-infections were performed with an equal number of mice infected with strains with their selection marker combinations reversed. For example, co-infections comparing wild type (Kan^R^) and mutant (Chlor^R^) were also performed with wild type (Chlor^R^) and mutant (Kan^R^). Bacterial titres from each organ and starting inoculum were used to calculate the competitive index (CI) as follows: CI = (Output wild type / Output mutant) / (Input wild type / Input mutant).

### RNA sequencing

Stationary phase cells were obtained by propagating strains statically for two serial 24 hour passages at 37°C, with a 1:1000 dilution between passages (identical to the inoculum used for mouse infections). Log phase cells were obtained by diluting overnight cultures 1:100 in LB and growing to log phase (OD_600_ = 0.4 – 0.5) at 37°C with shaking. RNA was extracted with the RNeasy Mini kit (Qiagen, Singapore) from three biological replicates for both log and stationary phase samples using 7 × 10^8^ cells. RNA quality was assessed using the Agilent RNA 6000 pico kit on an Agilent 2100 Bioanalyzer (Agilent Technologies, USA); only replicates where all samples had an RNA integrity number (RIN) >= 7 were used. Ribosomal RNA (rRNA) depletion was done with the Ribo-Zero rRNA removal kit (Epicenter, USA) according to the manufacturer’s recommended protocol. Libraries were generated using the ScriptSeq v2 RNA-seq library preparation kit (Epicenter, USA) according to the manufacturer’s recommended protocol. Each uniquely indexed strand specific library was assessed for library size and amount using the Agilent DNA 1000 kit (Agilent Technologies, USA). After normalization and pooling, samples were sequenced on either the Illumina HiSeq 4000 or NextSeq sequencer with 2 × 151 bp or 2 × 76 bp reads (Illumina, USA).

Raw sequencing reads were mapped to their respective reference genomes: RefSeq accession GCF_000013265.1 for *E. coli* UTI89, GCF_000005845.2 for *E. coli* MG1655 and GCF_000007445.1 for *E. coli* CFT073 using BWA-MEM (version 0.7.10) with default parameters (101). HTseq was used to quantify sequencing reads mapping to predicted open reading frames (ORFs) (102). Ribosomal RNA (rRNA) and transfer RNA (tRNA) sequences (based on the corresponding Genbank RefSeq annotation) were filtered out of the data set. R (version 2.15.1) was used for differential expression analysis, using the edgeR package (103). Briefly, samples were normalized by TMM (trimmed median of means), common and tagwise dispersion factors were estimated using a negative binomial model, and then fold change values were calculated from these normalized counts. A false discovery rate (FDR) cutoff of <= 0.05 and a log_2_ fold change >= 1.5 were applied, resulting in the final set of differentially expressed genes.

### Pacific Biosciences single molecule real time (SMRT) sequencing

Genomic DNA was extracted from log phase bacterial cultures grown in LB at 37°C with shaking and quantified using a Qubit 2.0 Fluorometer (Life Technologies, USA) using the dsDNA HS kit. 5 μg of DNA was sheared to 10 kbp using a g-Tube (Covaris, USA), and a SMRTbell library was made with the SMRTbell template prep kit 1.0 (Pacific Biosciences, USA) according to the manufacturer’s instructions. Library quality and quantity were assessed using the Agilent DNA 12000 kit (Agilent Technologies, USA) and sequenced on the PacBio RS II sequencer (Pacific Biosciences, USA). Sequencing was performed using a single SMRTCell with P4-C2 enzyme chemistry using a 180 min movie. Reads were mapped back to the corresponding reference genome (as indicated under “RNA Sequencing”) and methylated motifs were identified using the “RS_Modification_and_motif_analysis” algorithm in SMRT Analysis suite v2.3 using default parameters. Bases with a coverage of at least 25X and a kinetic score of at least 60 (default values for the RS_Modification_and_motif_analysis protocol) were identified as being methylated.

### Quantitative RT-PCR

Samples were prepared as described under “RNA sequencing”, using the RNeasy Mini kit (Qiagen, Singapore). 1 μg of RNA was treated with DNase I, RNase-free (Thermo fisher scientific, USA) at 37°C for 1 hour to remove any residual genomic DNA. Next, 500 ng of RNA was used for cDNA synthesis with SuperScript II Reverse Transcriptase and Random hexamers (Invitrogen, USA) according to the manufacturer’s recommended protocol. Amplified cDNA was diluted 1:4 for all target genes and 1:400 for the *rrsA* internal control. Real-time PCR was performed with the KAPA SYBR FAST qPCR kit (KAPA Biosystems, USA) on a LightCycler 480 instrument (Roche, Singapore) with the following cycle: 95°C for 5 mins, followed by 40 cycles of 95°C for 30 seconds and 60°C for 30 seconds. Target-specific primers for qRT-PCR were designed using the IDT PrimerQuest tool (IDT, Singapore) and are listed in S7 Table. Relative fold change for target genes was calculated by the ΔΔCT method utilizing the 16S ribosomal gene *rrsA* as the internal control. Reverse transcriptase and negative controls were included in each run.

### Motility Assay

Motility assays were performed as described previously with slight modifications (104). Strains were grown to log phase (OD_600_ = 0.4 – 0.5) by sub-culturing overnight bacterial cultures 1:100 in LB at 37°C. Bacteria were harvested by centrifugation (3200 g, 10 min) and resuspended in sterile PBS to OD_600_ = 0.4. Sterile 0.25% LB agar plates were prepared and stabbed once with each strain using sterile toothpicks. The soft agar plate was then incubated at 37°C for 7 – 8 hours. Motility was calculated by measuring the diameter of the bacterial motile front. Distances were normalized to a wild type control included in each experiment.

### Biofilm Assay

A 96-well Crystal Violet biofilm assay was performed as previously described (105). Briefly, strains were grown to log phase (OD_600_ = 0.4 – 0.5) by sub-culturing overnight bacterial culture 1:100 in LB at 37°C and resuspended in sterile PBS to OD_600_ = 0.4. 96-well clear flat bottom poly vinyl chloride (PVC) plates (Product #01816049, Costar, Singapore) were seeded with 200 μl of sterile media (LB or YESCA) and 5 μl of the PBS suspension. PVC plates were incubated for the specified times at 26°C or 37°C in a humidified chamber. Plates were washed once with water and stained with Crystal Violet for 30 mins. Excess stain was removed by washing thrice with water. Residual stain was then dissolved in 200 μl of 50% ethanol, with care taken to avoid disturbing the biofilm. The amount of biofilm was quantified by measuring OD at 590nm using a Sunrise 96 well microplate absorbance reader (Tecan, Switzerland). The biofilm produced by each test strain was normalized to that of the corresponding wild type strain.

### Growth Curves

Growth curves for bacterial strains were measured using the Bioscreen C instrument (Bioscreen, Finland). Strains were grown to log phase (OD_600_ = 0.4 – 0.5) by sub-culturing overnight bacterial culture 1:100 in LB at 37°C and resuspended in sterile PBS to OD_600_ = 0.4. 5 μl of this normalized bacterial suspension was then inoculated into 145 μl of the desired growth media (LB (rich) or M9 (minimal)) in triplicates. Plates were incubated at 37°C and OD_600_ was measured every 15 mins for 20 hours.

### Biolog Phenotype Microarray (PM)

PM assays (72) were performed using PM plates 1 to 20 by the PM services group at Biolog, USA using their standard protocol for *E. coli*. Succinate was added as the carbon source for PM plates 3-20 and all experiments with UTI89 (a niacin auxotroph) included 1 μg/ml niacin. All experiments were done with biological duplicates. Briefly, PM plates contain: PM1-2 Carbon sources, PM3 Nitrogen sources, PM4 Phosphorous and Sulphur sources, PM5 Nutrient supplements, PM6-8 Peptide nitrogen sources, PM9 Osmolytes, PM10 pH, and PM11-20 Inhibitors for different metabolic pathways. The PM services group analyzed growth curves, as measured colorimetrically, using their own proprietary software. Gain or loss of a phenotype/resistance was called by the PM services group, again using their proprietary software, which identified phenotypic differences based on the presence of a height difference between the strains in both replicates and a quality score >150 as cutoffs.

## Acknowledgements

The authors thank members of the Chen lab, Shu Sin Chng, William Burkholder, and Grishma Rane for helpful discussions and comments on the manuscript. The authors thank Steven Turner, Jonas Korlach, and Tyson Clark from Pacific Biosciences and Jacqueline L.Y. Chee for assisting with analysis of UTI89 methylation specificities using SMRT sequencing. The authors would also like to thank Adriano Baćac for additional technical assistance. Finally, the authors wish to thank the Next Generation Sequencing Platform at the Genome Institute of Singapore for technical help and guidance designing sequencing experiments.

## Authors’ contributions

KSM performed the experiments. KSM and SLC designed the experiments, analyzed the data, and wrote the manuscript. Both authors read and approved the final manuscript.

## Availability of data and materials

The PacBio sequencing data is available in the GenBank Short Read Archive under BioProject PRJNA474982 (106). The RNA-seq datasets generated and analysed during the current study are available in the GenBank Short Read Archive under BioProject PRJNA675406.

## Supporting information

**S1 Table. Methylated motifs identified by SMRT sequencing in *E. coli* UTI89 and UTI89 *ΔhsdSMR.*** Modified positions are underlined. DNA modifications: N6-methyladenine (6mA) and C5-methylcytosine (5mC). N=A, T, G or C and W=A or T. **\*** low percentage or absence of 5mC bases is due to the weak signal-to-noise ratio for this modification, requiring higher coverage or chemical conversion prior to sequencing (107,108).

**S2 Table. Logarithmic phase differentially expressed genes by RNA-seq.** Genes with log_10_ FDR <=0.05 and log_2_ fold change =>1.5 are considered significant and highlighted in bold. *denotes deleted genes/alleles and polar effects.

**S3 Table. Stationary phase differentially expressed genes by RNA-seq.** Genes with log_10_ FDR <=0.05 and log_2_ fold change =>1.5 are considered significant and highlighted in bold. *denotes deleted genes/alleles and polar effects.

**S4 Table. Phenotypic differences identified by PM due to altered *E. coli* Type I methylation**. Type I methylation mutants are compared to corresponding wt strains and phenotypic differences listed in descending value of quality score. Phenotypes with quality score >150 are considered significant and highlighted in bold. * denotes resistance conferred by presence of kanamycin selection cassette at *hsdSMR* locus.

**S5 Table. Distribution of Type I methylation motifs in***E. coli* **UTI89, MG1655 and CFT073 genomes.**\*UTI89 genome includes the 5,065,741bp chromosome and 114,230bp plasmid (pUTI89).

**S6 Table. List of strains and plasmids.**

**S7 Table. List of primers**.

**S1 Fig. Loss of native Type I methylation does not alter UTI89 urovirulence.** Wild type UTI89 or isogenic UTI89*ΔhsdSMR* strains were transurethrally inoculated (2 × 10^7^ cfu/mouse). Bladder and kidney pairs (as indicated by the labels below the x-axis) were aseptically harvested at 1 day post-infection (dpi), homogenized and plated for determination of bacterial burden (colony forming units (cfu)/ml). Mann-Whitney test was used to identify significant differences between strains in bladders and kidneys; * p <0.05, n = 5 mice/strain. Each data point represents a single mouse and horizontal lines represent the median values.

**S2 Fig. Deletion of UTI89 Type I RMS does not affect stationary phase gene expression.** RNA sequencing comparing stationary phase transcriptomes of UTI89 and UTI89*ΔhsdSMR.* Left, a volcano plot of log FDR against log fold change. Right, qq-plots showing the distribution of uncorrected p-values. Significantly differentially expressed genes (log_2_ fold change >1.5 and log_10_ FDR <0.05) are labelled and colored either red (deleted genes/polar effects) or blue (validated as false positive by qRT-PCR). n = 3 biological replicates.

**S3 Fig.***E. coli* **strains MG1655 and CFT073 encode functional archetypal Type I RMSs with distinct specificities. (A)** Transformation efficiency assay using plasmids bearing 0, 1, or 2 copies (as indicated on the x-axis) of the MG1655 Type I RMS motif (5’-AAC(N_6_)GTGC-3’). Recipient cells were wild type MG1655 and the isogenic *ΔhsdSMR* mutant, as indicated by the labels below the x-axis. **(B)** Transformation efficiency assay using plasmids bearing 0, 1, or 2 copies (as indicated on the x-axis) of the CFT073 Type I RMS motif (5’-GAG(N_7_)GTCA-3’). Recipient cells were wild type CFT073 and the isogenic *ΔhsdSMR* mutant, as indicated by the labels below the x-axis. Unmethylated and methylated plasmid preparations were used to transform each strain, as indicated by the legend at the top right. An unpaired t-test was used to identify significant differences between plasmids with 0, 1 and 2 methylation sites for both preparations and strains; * p <0.05, n = 3 biological replicates. Data represents mean ± s.d. of log transformed values.

**S4 Fig. Loss of Type I methylation in MG1655 and CFT073 has no effect on stationary phase gene expression or growth phenotypes. (A and B)** RNA sequencing comparing stationary phase transcriptomes of **(A)** MG1655 and MG1655*ΔhsdSMR*, and **(B)** CFT073 and CFT073*ΔhsdSMR.* Left, a volcano plot of log FDR against log fold change. Right, qq-plots showing the distribution of uncorrected p-values. Significantly differentially expressed genes (log_2_ fold change >1.5 and log_10_ FDR <0.05) are labelled and colored either red (deleted genes/polar effects) or brown (differentially expressed genes). n = 3 biological replicates. **(C and D)** Phenotype microarray (PM) panel with plates PM1 to 20 comparing **(C)** wt MG1655 and MG1655*ΔhsdSMR*, and **(D)** CFT073 and CFT073*ΔhsdSMR.* Each plate is represented as a 12×8 grid of growth curves (red (wt), green (*ΔhsdSMR*), and yellow (overlap) on a gray background). Each growth curve plots growth (measured colorimetrically) (y-axis) against time (x-axis). Wells representing conditions where a height difference was observed between the strains in both replicates are boxed in black, and wells which also have a quality score >150 are considered significant and boxed in red. n = 2 biological replicates.

**S5 Fig. UTI89 methylation-switch strain transcriptomes are unaffected by changing Type I methylation.** RNA sequencing comparing transcriptomes of corresponding growth phase wt UTI89 and **(A)** stationary phase UTI89 *hsdS^MG1655^*, **(B)** stationary phase UTI89 *hsdSMR^CFT073^*, **(C)** stationary phase UTI89 *hsdS^UTI89^*, and **(D)** log phase UTI89 *hsdS^UTI89^* Left, a volcano plot of log FDR against log fold change. Right, qq-plots showing the distribution of uncorrected p-values. Significantly differentially expressed genes (log_2_ fold change >1.5 and log_10_ FDR <0.05) are labelled and colored either red (deleted genes/polar effects) or brown (differentially expressed genes). n = 3 biological replicates.

**S6 Fig. UTI89 Type I methylation-switch does not alter growth phenotypes.** Phenotype microarray (PM) panel with plates PM1 to 20 comparing **(A)** wt UTI89 and UTI89 *hsdS^MG1655^*, and **(B)** UTI89 and UTI89 *hsdS^UTI89^.* Each plate is represented as a 12×8 grid of growth curves (red (wt), green (*ΔhsdSMR*), and yellow (overlap) on a gray background). Each growth curve plots growth (measured colorimetrically) (y-axis) against time (x-axis). Wells representing conditions where a height difference was observed between the strains in both replicates are boxed in black, and wells which also have a quality score >150 are considered significant and boxed in red. n = 2 biological replicates.

**S7 Fig.***E. coli* **growth is unaltered by Type I methylation.** Growth curves generated using optical density (OD_600_) values in **(A)** LB and **(B)** M9 minimal media. Wild type *E. coli* UTI89 (black), MG1655 (red), CFT073 (green), and corresponding Type I methylation mutant strains were tested, as indicated by the legend at the top right. Measurements were taken every 15 minutes and cultures propagated at 37℃. n = 3 biological replicates. Data points represent median values.

**S8 Fig. Type I methylation does not affect UTI89 motility.** Soft agar assay used to measure motility of UTI89 methylation mutant strains, as indicated by the legend at the top right. Motility is represented relative to wild type UTI89 motility, which was included in each replicate. Strain UTI89*ΔfliC* lacking flagellin serves as a non-motile control. Wilcoxon signed rank test was used to identify significant differences in bacterial motility relative to wild type; * p <0.05, n = 3 biological replicates. Data represents median and 95% confidence intervals.

**S9 Fig. UTI89 biofilm formation is not affected by Type I methylation.** 96-well crystal violet assay for biofilm quantification using LB **(A and B)** or YESCA **(C and D)** media. UTI89 Type I methylation mutant strains were tested, as indicated by the legend at the top right. Assay was performed for 24 hours **(A and C)** or 48 hours **(B and D)**, as well as at 26℃ and 37℃ for each media as indicated by the labels below the x-axis. Biofilms were quantified using crystal violet absorbance at 590nm (OD_590_) and represented relative to wild type UTI89 which was included in each replicate and condition. Wilcoxon signed rank test was used to identify significant differences in biofilm formation relative to wild type; * p <0.05, n = 3 biological replicates. Data represents median and 95% confidence intervals.

**S10 Fig. MG1655 Type I methylation-switch has no effect on growth phenotypes.** Phenotype microarray (PM) panel with plates PM1 to 20 comparing **(A)** wt MG1655 and MG1655 *hsdS^UTI89^*, and **(B)** MG1655 and MG1655 *hsdS^MG1655^.* Each plate is represented as a 12×8 grid of growth curves (red (wt), green (*ΔhsdSMR*), and yellow (overlap) on a gray background). Each growth curve plots growth (measured colorimetrically) (y-axis) against time (x-axis). Wells representing conditions where a height difference was observed between the strains in both replicates are boxed in black, and wells which also have a quality score >150 are considered significant and boxed in red. n = 2 biological replicates.

